# Machine Learning-Assisted Digital PCR and Melt Enables Broad Bacteria Identification and Pheno-Molecular Antimicrobial Susceptibility Test

**DOI:** 10.1101/587543

**Authors:** Pornpat Athamanolap, Kuangwen Hsieh, Christine M. O’Keefe, Ye Zhang, Samuel Yang, Tza-Huei Wang

## Abstract

Toward combating infectious diseases caused by pathogenic bacteria, there remains an unmet need for diagnostic tools that can broadly identify the causative bacteria and determine their antimicrobial susceptibilities from complex and even polymicrobial samples in a timely manner. To address this need, a microfluidic- and machine learning-based platform that performs broad bacteria identification (ID) and rapid yet reliable antimicrobial susceptibility testing (AST) is developed. Specifically, this new platform builds on “pheno-molecular AST”, a strategy that transforms nucleic acid amplification tests (NAATs) into phenotypic AST through quantitative detection of bacterial genomic replication, and utilizes digital PCR and digital high-resolution melt (HRM) to quantify and identify bacterial DNA molecules. Bacterial species are identified using integrated experiment-machine learning algorithm via HRM profiles. Digital DNA quantification allows for rapid growth measurement that reflects susceptibility profiles of each bacterial species within only 30 min of antibiotic exposure. As a demonstration, multiple bacterial species and their susceptibility profiles in polymicrobial urine specimen were correctly identified with a total turnaround time of ~4 hours. With further development and clinical validation, this new platform holds the potential for improving clinical diagnostics and enabling targeted antibiotic treatments.

Table of Contents Graphic

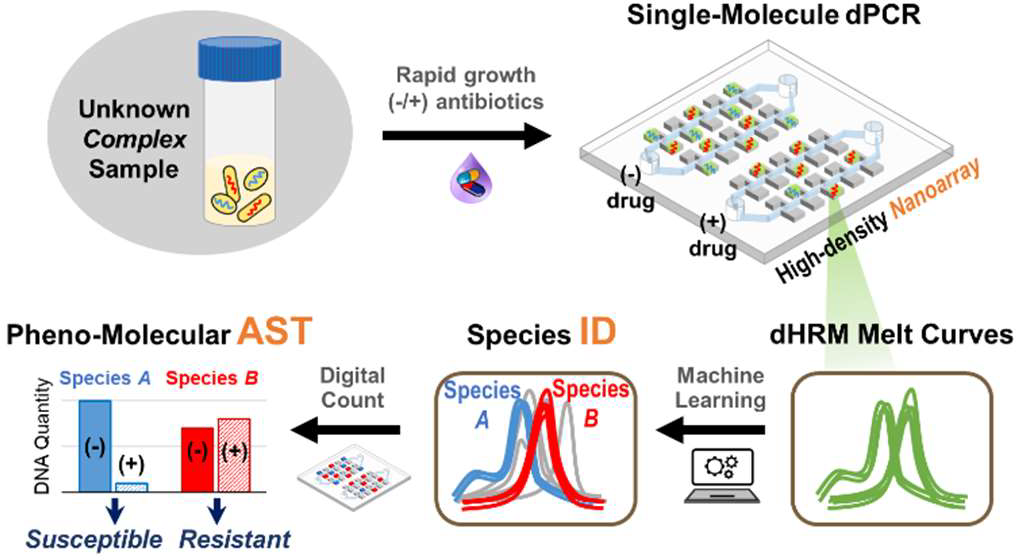

## Introduction

Infectious diseases inflicted by pathogenic bacteria continue to threaten human health and incur heavy economical burdens. For example, sepsis is the leading cause of deaths in hospitals^[1,2]^ and approximately 270,000 Americans die from sepsis each year.^[3]^ Similarly, urinary tract infections (UTIs) affect approximately 50% of women at least once in their lifetime and incur > $2 billion in treatment cost in the United States per year.^[4–8]^ Adding severity to the problem, recent reports reveal that these bacterial infections (*e.g*. UTIs^[9–12]^ and wound infections^[13–15]^) can be polymicrobial, often leading to increased infection severity and poorer patient outcome.^[16–21]^ To combat this serious threat, tools for diagnosing infectious diseases must be improved. A comprehensive clinical diagnosis must encompass broad identification (ID) of the causative bacteria and antimicrobial susceptibility test (AST). Critically, the diagnosis must be made rapidly (*e.g*., within a few hours) and ideally from various sample matrices of different diseases. Unfortunately, traditional diagnostic methods cannot meet these requirements as they rely heavily on bulk bacterial culture that can take several days or even up to weeks to complete.^[22]^ The lengthy lag to definitive diagnosis obtained via traditional diagnostic methods results in the common use of broad-spectrum antibiotics, which can lead to poor patient outcome and rampant spread of antimicrobial resistance.^[23,24]^ Consequently, there remains a clear need for new diagnostic technologies that can meet the multi-faceted requirements in the diagnosis of infectious diseases.^[25]^

By detecting specific genes within bacteria with high sensitivity, specificity, and speed, nucleic acid amplification tests (NAATs) such as PCR provide an effective foundation for accelerating bacteria ID and, to some extent, AST from days to hours. Employing NAATs for bacteria ID is now a turnkey process and broadly falls under two approaches. In the first approach, a panel of bacterial species-specific primers or probes is employed in a multiplexed assay to identify specific bacteria within the panel. Several FDA-approved commercial platforms (most notably FilmArray from BioFire Diagnostics, BD Max from Becton Dickinson Diagnostics, and GeneXpert from Cepheid) utilize this multiplexed detection strategy to identify bacteria. Though efficient, this approach would yield false negative tests for pathogenic bacteria outside of the existing panel. In the other approach, universal PCR using a pair of pan-bacteria primers is first performed to detect any bacterial species, followed by post-amplification analysis techniques such as high-resolution melt (HRM) to achieve species identification. Although requiring prior efforts in building the database of molecular profiles (*e.g*., melt curves) of bacteria, HRM provides a simple, yet practical post-PCR sequence fingerprinting solution for large-scale bacterial ID.^[26–32]^ Employing NAATs for AST, in contrast to the maturity of bacteria ID, remains a work in progress. NAATs can detect genetic markers such as mutations that confer antibiotic resistance, which can serve as a surrogate for determining antibiotic susceptibility. Unfortunately, except for a few well-established markers (*e.g*., mecA^[33–35]^ and vanA and vanB^[36,37]^), resistance genes cannot reliably predict susceptibility to a particular antibiotic.^[38,39]^ Therefore, advanced NAAT-based methods that can identify bacteria and provide reliable AST information must be developed.

“Pheno-molecular AST” is an emerging approach that achieves reliable AST and concurrent bacteria detection by combining phenotypic characterization of antibiotic susceptibility with quantitative, nucleic-acids-based molecular detection of bacteria.^[22,40–50]^ In pheno-molecular AST, bacteria are first briefly incubated in the presence and absence of antibiotics. The amounts of bacterial nucleic acids – serving as surrogates of bacterial growths – between antibiotic-treated samples and no-antibiotic controls are then quantitatively detected and compared to determine the antibiotic susceptibilities. Moreover, when coupled with NAATs for quantitative detection, pheno-molecular AST can significantly outpace traditional culture-based AST in sensitivity and assay time, while adding a layer of specificity. To date, several pheno-molecular AST assays based on real-time PCR^[40]^, digital PCR^[45]^, and digital LAMP^[46]^ have been demonstrated. Unfortunately, no platform has reported pheno-molecular AST with broad bacteria ID in a digital format for achieving a comprehensive and rapid diagnostic method. For example, Chen et al.^[40]^ reported a pheno-molecular AST assay that specifically targeted *N. gonorrhoeae*. Schoepp et al.^[45,46]^ developed microfluidic-based, digital pheno-molecular AST assays that shortened the antibiotic exposure time to 15 to 30 min, though they only focused on *E. coli* as the target organism. Meanwhile, although we and others^[47,48]^ have combined pheno-molecular AST with real-time, quantitative universal PCR and HRM analysis to achieve broad identification, these bulk-based assays have a limited ability of analyzing polymicrobial samples because composite melt curves from multiple bacteria species cannot be easily decoupled and resolved.

In response, we have developed the first universal digital PCR and HRM (dPCR-HRM) platform for performing broad bacteria ID and rapid pheno-molecular AST toward clinical diagnosis of infectious diseases. At the center of our platform is the Nanoarray – a microfluidic device that we have engineered to perform dPCR-HRM upon digitizing single bacterial DNA molecules. In doing so, our Nanoarray not only facilitates precise quantification of bacterial DNA molecules but also ensures that each melt curve in the device is generated from a single bacterial species, which allows us to identify individual bacterial species even from polymicrobial samples. We have also developed a machine learning-based algorithm for the analysis of digital HRM (dHRM) profile from each bacterial DNA to achieve species identification and accurate quantification. Antibiotic susceptibility of each species can thus be determined based on digital counts of DNA quantity that reflects bacterial growth under antibiotic exposure. The quantitative precision of our dPCR-HRM platform allows for rapid antibiotic exposure time in as little as 30 min. As an additional benefit, our method also works with bacteria in urine, a complex sample matrix. For demonstration, we used our method to correctly identify gentamicin-sensitive *E. coli* and gentamicin-resistant *Staphylococcus aureus* that were both present in a polymicrobial urine sample with a total turnaround time of ~4 hours, illustrating the potential of our method toward clinical diagnosis.

## Results and Discussion

### Assay Overview

We have developed a streamlined, dPCR-HRM-based workflow for performing broad bacteria ID and pheno-molecular AST from complex samples. We begin by evenly dividing the sample into two aliquots, exposing one aliquot to an antibiotic and the other to no antibiotics (*i.e*., no-drug control), and incubating both aliquots at 37 °C for 30 min (Figure 1, Step 1). Following incubation, bacterial DNA from both aliquots are extracted in parallel, diluted (if necessary), mixed with universal PCR mixture, and loaded into two independent modules on a Nanoarray device comprising of thousands of nanoliter reaction wells (nanowells) (Figure 1, Step 2). The use of limiting dilutions with Nanoarrays leads to digitization of DNA molecules in the nanowells for dPCR and dHRM and that each melt curve is generated from a single bacterial species. In doing so, even multiple species in heterogeneous samples can be individually and independently enumerated. Next, we place the device onto our custom thermal-optical platform^[51]^ (Figure S1), which performs thermocycling for dPCR and temperature ramping for dHRM, and acquires fluorescence images of the entire Nanoarray at all temperature increments during dHRM. These “temperature-lapse” fluorescence images are then compiled to generate digital melt curves for all nanowells within the device in parallel. Each digital melt curve in a nanowell is analyzed via our in-house, machine-learning-based melt curve identification algorithm to match to the species-specific melt curves in our digital melt curve database, thereby identifying the bacterial species present in the nanowell (Figure 1, Step 3-ID). The total number of bacterial species-specific DNA molecules from each aliquot can then be accurately counted. Finally, antibiotic susceptibility of each bacterial species is determined by comparing the number of DNA molecules of the drug-treated aliquot with the no-drug control aliquot, where significantly fewer DNA counts indicate susceptibility to the antibiotic while comparable DNA counts indicate resistance to the antibiotic (Figure 1, Step 3-AST).

**Figure 1.**
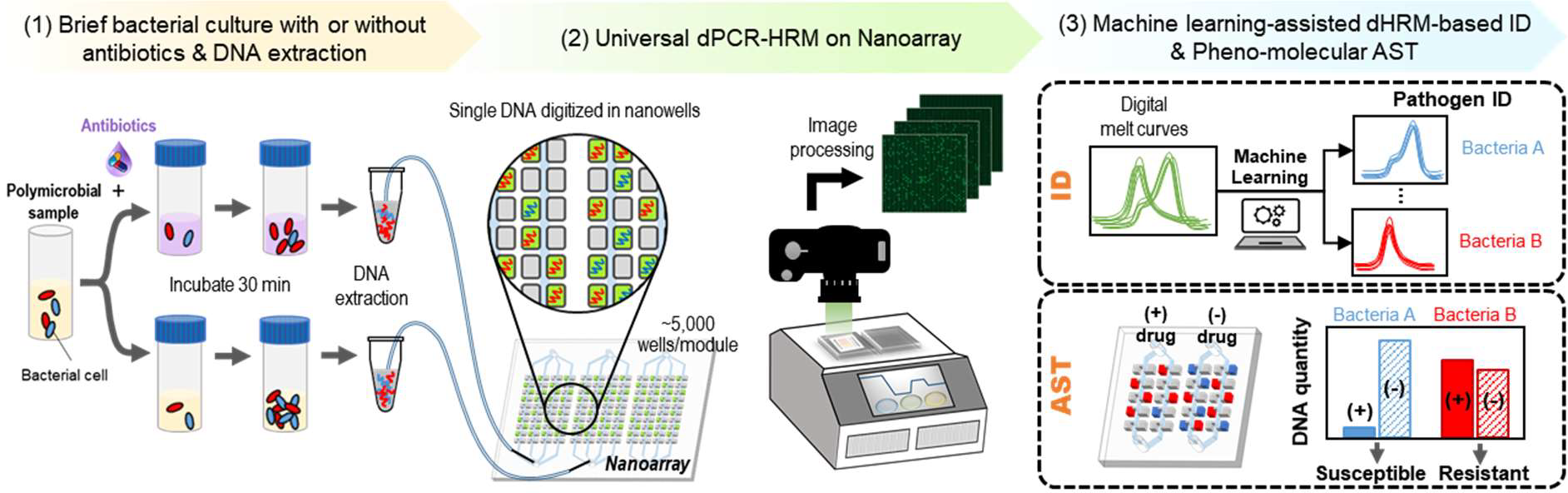
Overview of digital PCR and melt platform with machine learning-assisted algorithm for rapid bacteria ID and pheno-molecular AST. Analyzing bacterial growth under antibiotic exposure by measuring bacterial genome replication (*i.e.*, pheno-molecular AST) via universal digital PCR and high resolution melt (dPCR-HRM) in the Nanoarray offers an effective means for achieving rapid yet comprehensive bacteria ID and AST, even from complex samples (*e.g*., urine) that contain multiple bacteria with distinct antibiotic susceptibilities. The streamlined workflow begins with (1) briefly incubating evenly divided bacterial sample aliquots with and without antibiotics and extracting bacterial DNA from both aliquots. Next, (2) bacterial DNA from each aliquot is mixed with a universal PCR mixture that contain pan-bacteria primers and Evagreen dye, loaded into separate modules of a Nanoarray device, and placed on a thermal-optical platform to perform dPCR-HRM. The ~5,000 nanoliter reaction wells (*i.e.*, nanowells) in each module of the Nanoarray, along with performing limiting dilution of both aliquots, ensure that only one DNA molecule is digitized and subsequently PCR-amplified in each nanowell, which guarantees that each melt curve is generated from a single bacterial species. During dHRM, fluorescence images of the entire Nanoarray are captured by the thermal-optical platform at specified temperature increments. (3) After dPCR-HRM, fluorescence intensities within all nanowells are extracted from these “temperature-lapse” fluorescence images to generate digital melt curves for all nanowells. The digital melt curves are analyzed by a machine learning-based melt curve identification algorithm to identify the bacterial species. Next, the number of species-specific digital melt curves is enumerated to quantify the DNA copy number for each bacterial species in each aliquot. Finally, the comparison between the DNA copy numbers of the two aliquots reveals the antibiotic susceptibility – higher DNA copies from the no-antibiotic aliquot indicate susceptibility while comparable DNA copies from both aliquots indicate resistance.

### Validation of dPCR-HRM in Nanoarray and Construction of Digital Melt Curve Database

An enabling element of our platform is the Nanoarray – a microfluidic device that we have engineered to rapidly and efficiently digitize single bacterial DNA molecules and reliably perform dPCR-HRM. In the current iteration, our Nanoarray features 3 identical but independent modules; each of these high-density modules houses 5,040 1-nL nanowells (Figure 2A). The 3 parallel modules within a single device allow us to analyze both drug-treated and no-drug AST conditions while performing a control dPCR-HRM with a well-characterized DNA sample. To ensure efficient and reliable digitization of bacterial DNA molecules, as well as robust dPCR-HRM, the Nanoarray was fabricated with a thin PDMS layer (~100 µm) sandwiched between a top glass coverslip and a bottom glass slide (Figure S2). Air-permeable PDMS in the Nanoarray facilitates vacuum-assisted DNA sample loading^[51,52]^, which allows the sample to fill all nanowells in a pre-desiccated device in < 5 s. A brief injection of a partitioning oil into the device is then sufficient for isolating all sample-filled nanowells. During dPCR, the combination of the thin PDMS layer and the top glass coverslip minimizes evaporation of nanoliter-scale reactions within nanowells. The bottom glass slide, which is 1 mm in thickness, creates a rigid and flat bottom surface of the device that ensures the temperature uniformity across the device during dPCR-HRM. Finally, we note that the partitioning oil contains PDMS, which solidifies during thermocycling to permanently encapsulate PCR products in each nanowell, thereby preventing cross-contamination and easing any device handling during dHRM analysis.

**Figure 2.**
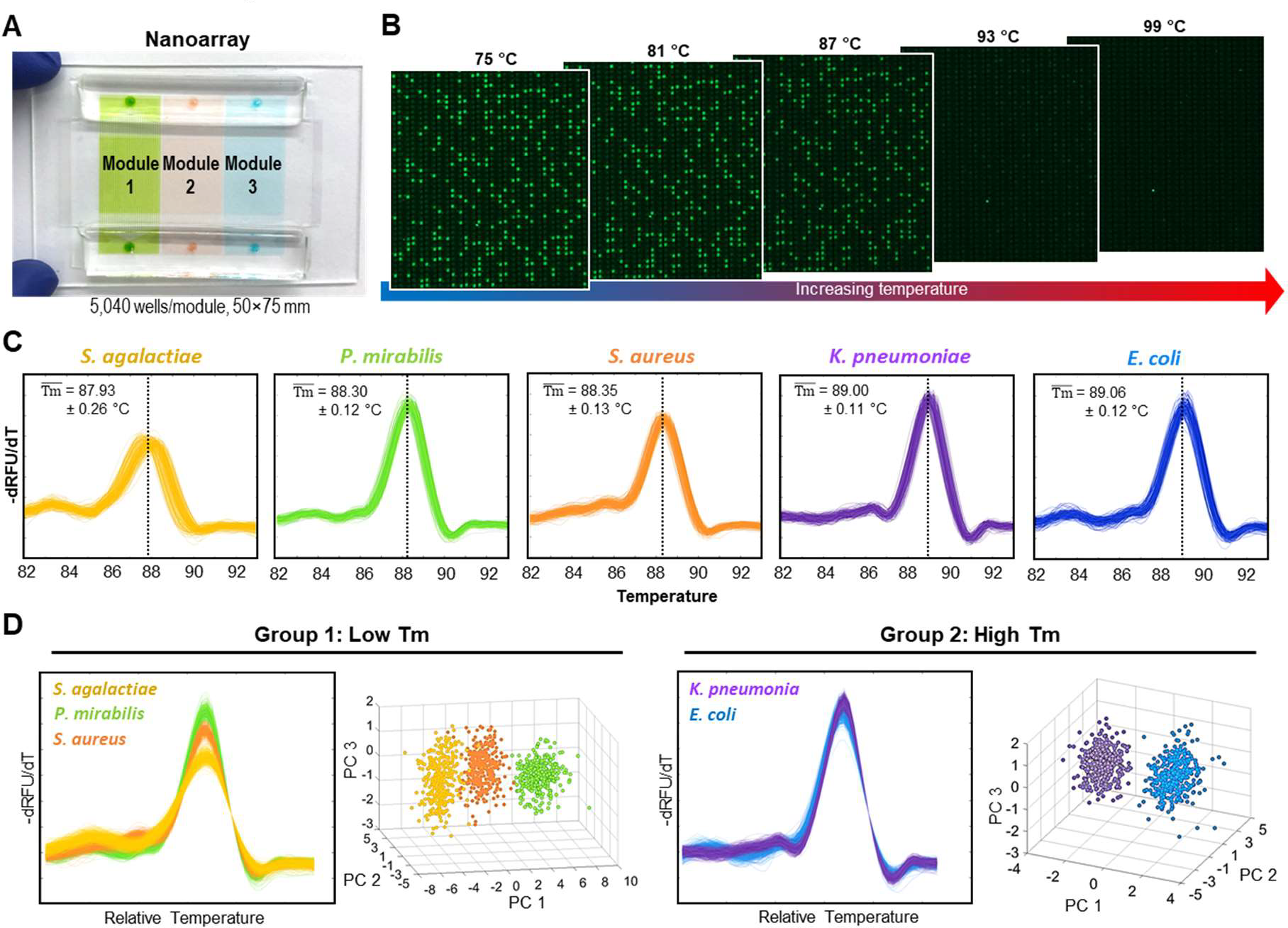
Bacteria ID via dPCR-HRM in Nanoarray and machine learning-assisted digital melt curve identification. A) Each Nanoarray device contains 3 independent modules for analyzing two samples and including a dPCR-HRM control. Each module houses 5,040 nanowells that are 1-nL in volume. B) Upon the completion of dPCR and immediately before the commencement of dHRM, strong green fluorescence can be observed from nanowells that have digitized and PCR-amplified bacterial DNA. During dHRM, double-stranded dPCR products in these strongly fluorescent, positive wells become increasingly melted as temperature increases, resulting in decreasing fluorescence intensities in these nanowells. Parallel measurements of fluorescence intensities in thousands of nanowells as a function of temperature lead to thousands of digital melt curves from a single experiment in a Nanoarray. C) 320 digital melt curves from 5 species of bacteria commonly found in urinary tract infections – *S. agalactiae*, *P. mirabilis*, *S. aureus*, *K. pneumoniae*, and *E. coli* – are collected to build a digital melt curve database toward broad bacteria ID. D) To achieve reliable bacteria ID, both the melting temperature (*Tm*) and the shape of the digital melt curves are used for analysis. Based on *Tm*, our digital melt curve database is divided into the low *Tm* group with *S. agalactiae*, *P. mirabilis*, and *S. aureus*, and the high *Tm* group with *K. pneumoniae*, and *E. coli*. Within each *Tm* group, digital melt curves are aligned to a single point to facilitate shape-based digital melt curve analysis before one-versus-one support vector machine algorithm is used to compare species-specific melt curve shapes and identify bacterial species. Also within each *Tm* group, principle component analysis using the first three principle components (*i.e*., PC1, PC2, and PC3) is performed to visualize that the digital melt curves from each species indeed cluster into distinguishable populations, thus confirming their distinct shapes.

Within the Nanoarray, we have implemented a universal dPCR-HRM assay that can amplify a broad range of bacterial DNA in the sample and subsequently differentiate and identify the bacterial species. This is achived by employing a pair of universal PCR primers that hybridize to conserved regions (*i.e*., consistent among all bacteria) flanking a hypervariable region (*i.e*., unique to each bacterial species) in the 16S rRNA gene and Evagreen dye that facilitates HRM analysis. We first validated our universal dPCR-HRM assay in the Nanoarry using *E. coli* genomic DNA as the target. In this initial validation, we loaded the target DNA and PCR mixture into a module of a Nanoarray device at a “digital concentration” such that only some of nanowells would be filled with a single copy of *E. coli* DNA. After dPCR, we indeed observed many “negative” nanowells that showed weak green fluorescence signal comparable to only surrounding background and channels, indicating that no dPCR occurred in these nanowells as they contained no target DNA (Figure 2B, left). Importantly, we also detected a number of “positive” nanowells that exhibited strong green fluorescence in the module, suggesting that a single copy of the target DNA was digitized and amplified in each of these nanowells. Subsequently during dHRM, temperature-lapse fluorescence images of the entire module revealed that fluorescence signals in all positive nanowells decreased as the temperature ramped up, indicating that DNA amplicons in these nanowells became increasingly melted (Figure 2B). After extracting fluorescence intensities within nanowells from the temperature-lapse fluorescence images, we obtained hundreds high-resolution digital melt curves for *E. coli* (Figure 2C, *E. coli*, blue). The digital melt curves not only closely resembled each other, but also the melt curve of *E. coli* obtained from a benchtop PCR-HRM (Figure S3). These results provide strong validation for our universal dPCR-HRM in the Nanoarray.

We then performed dPCR-HRM in our Nanoarray to detect and generate digital melt curves for 4 additional bacteria, thereby building a digital melt curve database with 5 common UTI bacterial species toward bacteria identification. As expected, our universal dPCR-HRM successfully detected genomic DNA from *Streptococcus agalactiae* (ATCC 13813), *Proteus mirabilis* (ATCC 12453), *S. aureus* (ATCC 29213), and *Klebsiella pneumonia* (ATCC BAA-1705) (Figure S4) and produced hundreds of digital melt curves for each species (Figure 2C). For all 5 species, the digital melt curves universally exhibited the single main peak profile – an indication that the melt curves were derived from single species. Importantly, however, the digital melt curves of each species differed from those of the other 4 species in their melting temperature (*Tm*) and their shape. For example, the digital melt curves from *E. coli* (Figure 2C, blue; *Tm* = 89.06 ± 0.12 °C) and *P. mirabilis* (Figure 2C, green; *Tm* = 88.30 ± 0.12 °C) showed similar shapes but differed by nearly 0.8 °C in average *Tm*. The digital melt curves from *S. aureus* (Figure 2C, orange; *Tm* = 88.35 ± 0.13 °C) displayed a distinctive ramp leading to the main peak. The digital melt curves from *S. agalactiae* (Figure 2C, yellow; *Tm* = 87.93 ± 0.26 °C) exhibited a small bump before a low main peak. Finally, the digital melt curves from *K. pneumoniae* (Figure 2C, purple; *Tm* = 89.00 ± 0.11 °C) had a small but noticeable drop immediately before the main peak. These differences form the basis for achieving bacteria ID based on their unique melt curves.

### Machine Learning-Assisted Algorithm for Digital Melt Curve and Bacteria ID

Having built our digital melt curve database, we next developed a new identification algorithm for distinguishing and identifying these species-specific digital melt curves. Our algorithm includes four key steps: 1) incorporation of *in situ* reference digital melt curves from *E. coli*, 2) classification of digital melt curves into relative *Tm*-based groups, 3) normalization and alignment of digital melt curves, and 4) identification of digital melt curves based on their shapes using a machine learning-based algorithm. In our platform, in addition to performing dPCR-HRM for the two samples of interest in two modules of the Nanoarray, we concurrently perform dPCR-HRM with purified genomic DNA of *E. coli* in the third module to generate reference digital melt curves. We then use relative *Tm* (*i.e*., differences in *Tm* between the digital melt curves of interest and the reference digital melt curves) to classify the digital melt curves of interest into groups that contain a subset of the bacterial species within the database (Figure S5). This relative *Tm*-based classificaition strategy is more consistent than directly using the exact *Tm* of the digital melt curves of interest toward identification, as the *Tm* can vary from experiment to experiment due to variations in devices and the non-uniform temperature distribution of the heating instrument. Following relative *Tm*-based classification, to enable the identification of digital melt curves of interest based on their shapes, we normalize the area under the curve of all digital melt curves and then align them to a single point. Finally, using an adapted in-house developed digital melt curve classification tool based on one-versus-one support vector machine (ovoSVM) algorithm^[26,48,53]^, the shape of each digital melt curve of interest is compared to the digital melt curves in the *Tm*-classified group to identify the bacterial species represented by the digital melt curve of interest.

Using a total of 1,600 digital melt curves – 320 from each of the 5 bacterial species in our database – we illustrate the outcomes of the key steps and demonstrate the overall performance of our machine learning-assisted algorithm in bacteria identification. In our database, digital melt curves of *S. agalactiae*, *P. mirabilis*, and *S. aureus* have lower *Tm* than those of *E. coli*, while digital melt curves of *K. pneumoniae* have comparable *Tm* as those of *E. coli* (Figure 2C). Classification of these digital melt curves based on *Tm* therefore results in the low *Tm* group with *S. agalactiae*, *P. mirabilis*, and *S. aureus*, and the high *Tm* group with *K. pneumoniae* and *E. coli* (Figure 2D). After normalization and alignment of digital melt curves, we performed principal component analysis (using a built-in Matlab function) to show that the digital melt curves within each group have distinct shapes. Indeed, principle component analysis for the low *Tm* group results in three distinct clusters, indicating that *S. agalactiae*, *P. mirabilis*, and *S. aureus* indeed have distinctly-shaped digital melt curves (Figure 2D-left). Principle component analysis similarly reveals that the digital melt curves of *K. pneumoniae* and *E. coli* in the high *Tm* group have distinct shapes (Figure 2D-right). Notably, principle component analysis shows that, when put in a single group, the digital melt curves of the 5 species are not fully distinguished (Figure S6). The analysis therefore confirms the importance of *Tm*-based classification toward robust identification. Finally, we iteratively performed 320 leave-one-out cross-validation experiments to validate our identification algorithm. That is, for each bacterial species, we trained our algorithm using 319 digital melt curves and then challenged our algorithm to identify the last digital melt curve, and repeated until all 320 digital melt curves had been tested. Our identification algorithm correctly identified 98.5% of the 1600 digital melt curves, illustrating the effectiveness of our machine learning-assisted algorithm in the identification of digital melt curves, and hence, bacterial species.

### Bacterial DNA Quantification in Nanoarray

Precise quantification of DNA molecules is essential for pheno-molecular AST. To this end, our Nanoarray supports digital quantification of DNA molecules through both dPCR alone and dPCR-HRM. Although dPCR alone allows for precise quantification of DNA, our dPCR-HRM can offer comparable level of precision as dPCR while adding the dHRM-based verification step that can remove false-positives from the counting results. For example, when we performed a no template control (NTC) experiment, 6 nanowells (0.12%) showed strong fluorescence signal after dPCR and would be considered as positives. However, upon analysis with our identification algorithm, we found that 4 out of these 6 “positives” had digital melt curves that matched poorly with any of the digital melt curves from the 5 bacterial species in our database. These poorly matched digital melt curves likely arose from non-specific amplifications such as primer dimers and were thus relegated as false positives. The remaining 2 positives were identified as *E. coli* and *K. pneumoniae* that likely stemmed from bacterial DNA contamination that may exist even in DNA polymerase^[54,55]^ (Figure S7). This result demonstrates that dHRM analysis in Nanoarray enhances assay specificity, as false-positives from non-specific melt curves or contaminants can be revealed, examined, and potentially eliminated.

To demonstrate quantification of DNA molecules at concentrations relevant to our subsequent pheno-molecular AST in the Nanoarray, we performed dPCR-HRM for 3 dilutions of purified *E. coli* genomic DNA at estimated 0.01, 0.1, and 1 genomic copy per nanowell, and compared the target quantification results from dPCR alone and from dPCR-HRM. Based on dPCR alone, we counted 5.4%, 40.1%, and 99.3% positives for these 3 dilutions (Figure 3A and B). Using dPCR-HRM for analysis, we determined that a fraction of these “positives” in each dilution were non-*E. coli* and consequently modified the counts to 5.3%, 39.5%, and 99.2% positives for these 3 dilutions (Figure 3C). These results support that our Nanoarray facilitates precise quantitative detection of DNA molecules and that dPCR-HRM can further improve quantification accuracy.

**Figure 3.**
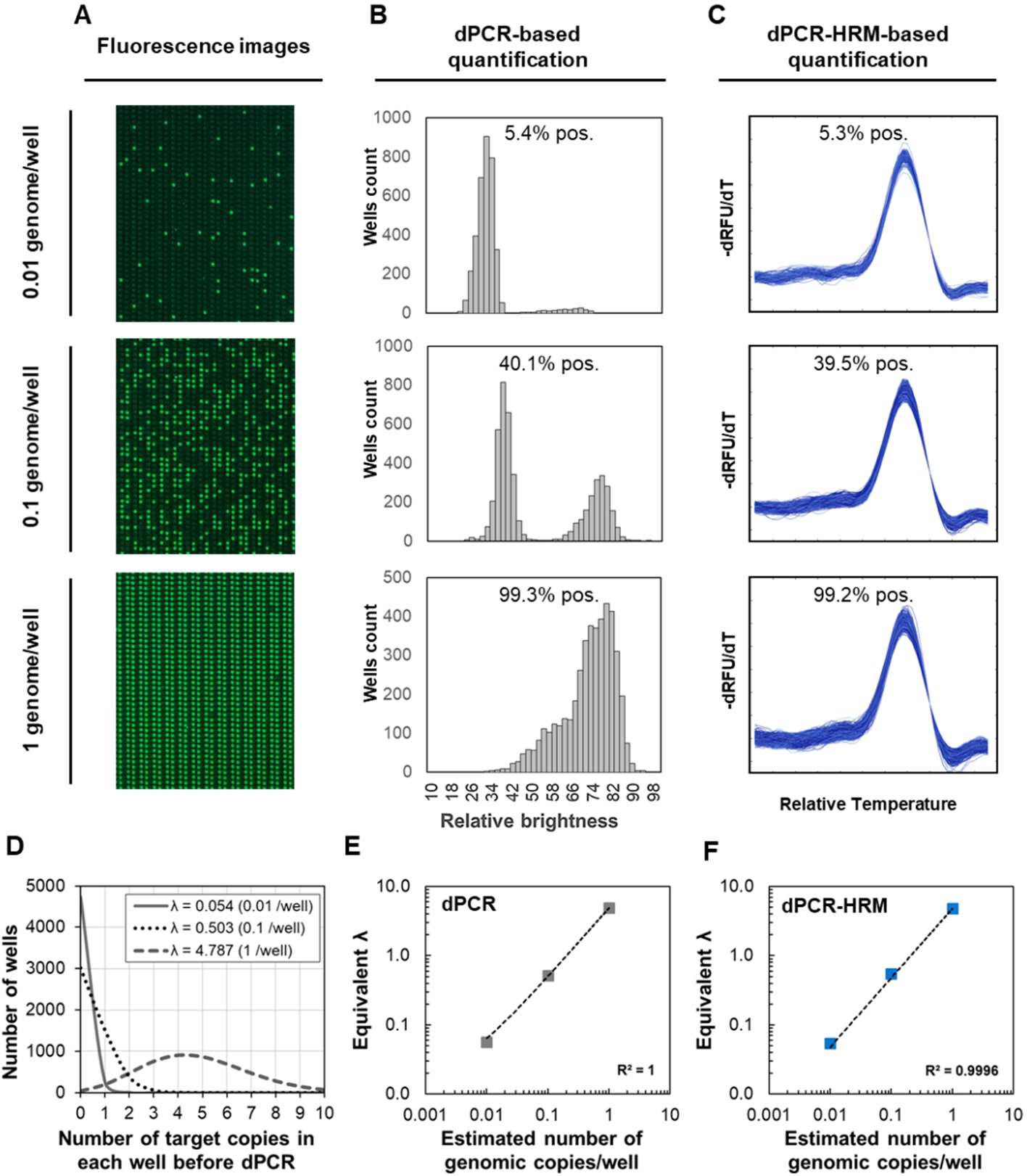
Digital bacterial DNA quantification in Nanoarray. The Nanoarray supports digital quantification of bacterial DNA through dPCR alone or dPCR-HRM. A) After performing dPCR for 3 concentrations of *E.coli* genomic DNA (estimated at 0.01, 0.1, and 1 copy per nanowell), increasing numbers of positive nanowells can indeed be observed from the fluorescence images of the 3 Nanoarray modules as the DNA concentration increases. B) In dPCR-based quantification, which is based only on the fluorescence level within nanowells, a histogram of relative fluorescence brightness is plotted to quantify the percentage of positive wells in each Nanoarray module for each DNA concentration. Based on dPCR, 5.4%, 40.1%, and 99.3% positive nanowells are detected for the 3 DNA concentrations. C) In contrast, dPCR-HRM-based quantification relies on counting the number of correctly-identified digital melt curves, which can enhance the assay specificity. Indeed, dPCR-HRM reveals that a fraction of the “positive nanowells” in each dilution originates not from *E. coli* but instead non-specific amplifications. The percentages of positive nanowells are therefore re-calculated to 5.3%, 39.5%, and 99.2% for these 3 dilutions. D) The percentage of positive nanowells from of each concentration can be used to calculate the corresponding mean occupancy (λ). Based on the calculated λʼs, the number of nanowells containing different copies of target in each nanowell can be visualized via Poisson distribution. A strong linear relationship between the estimated number of genomic copy per nanowell and λ can be observed from these 3 input concentrations quantified via both E) dPCR and F) dPCR-HRM, which provides strong evidence of precise quantification of DNA molecules in the Nanoarray and comparable precision in DNA quantification between dPCR and dPCR-HRM.

Consistent with standard digital-based quantification methods, we can calculate the quantity of our *E. coli* target based on the percent positive for each dilution. This calculation corrects for positive nanowells that contain more than one copy of the target. Specifically, we applied Poisson distribution to calculate the mean occupancy of DNA per nanowell, λ, using Equation 1

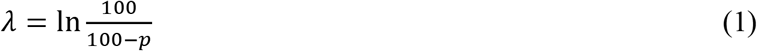

where *p* represents the percent positives. Hereafter, we used λ as our standard quantitation metric. Based on the percent positives from dPCR, the λʼs were 0.056, 0.513, and 4.906 for the 3 dilutions. Using the percent positives from dPCR-HRM, the λʼs were 0.054, 0.503, and 4.787 for the 3 dilutions. We can also calculate the probability distribution of the number of DNA molecules per nanowell *(k)* that would result from λ of each dilution calculated from our dPCR-HRM results using Equation 2 (Figure 3D, based a total of 5,000 wells).

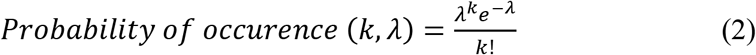

More importantly, we observed strong linear relationships between λʼs and the input genomic copy numbers for both dPCR (Figure 3E) and dPCR-HRM (Figure 3F). The excellent linearity in both cases provides strong support for precise quantification of DNA molecules in the Nanoarray and comparable precision in quantification between dPCR and dPCR-HRM. Finally, we note that a single *E. coli* genome contains ~7 copies of 16S rRNA gene^[56]^, which explains why we measured ~5 times more DNA than the estimated input from our 3 dilutions of *E. coli* target. The small difference is presumably due to typical DNA fragmentation during DNA extraction and locations of the 7 copies of the gene. Specifically, fragment sizes of extracted DNA generally range from 20 to 200 kb^[57,58]^, while the distance between copies of the 16S rRNA gene ranges from 40 to 2,700 kb.^[59]^ It is thus likely that *E. coli* genomic DNA molecules were fragmented such that most copies of the 16S rRNA gene were digitized into separated nanowells, while some adjacent 16S copies were in the same fragments, resulting in fewer positives that could be measured in the Nanoarray.

### dPCR-HRM-based Pheno-Molecular AST in Nanoarray

The capacity for digitally detecting even minute increases in the quantity of DNA molecules via dPCR-HRM in the Nanoarray enables brief antibiotic exposure and consequently rapid pheno-molecular AST. To demonstrate this concept, we incubated gentamicin-susceptible and gentamicin-resistant strains of *E. coli* with gentamicin (a commonly used intravenous antibiotic for infections caused by gram-negative bacteria such as *E. coli*) at either 0 or 4 μg mL^−1^ (the low end of the clinical breakpoint for resistance/susceptibility) for 30 min before quantifying the DNA molecules from each case via dPCR-HRM in the Nanoarray. For the gentamicin-susceptible *E. coli* strain, λ of 0.120 (*i.e*., 0.120 λ) and significantly lower 0.042 λ were measured for the no-gentamicin control and the gentamicin-treated sample, respectively, suggesting that gentamicin indeed inhibited the growth of this susceptible *E. coli* (Figure 4A). Conversely, for the gentamicin-resistant *E. coli* strain, comparable 0.037 λ and 0.033 λ were observed from the no-gentamicin control and the gentamicin-treated sample, respectively, indicating that gentamicin could not inhibit the growth of this resistant *E. coli* (Figure 4B). By taking the ratio between λ of the drug-treated sample and λ of the no-drug control (*i.e*., Equation 3),

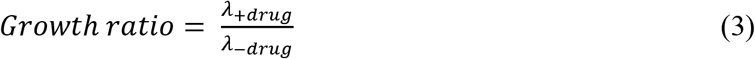

**Figure 4.**
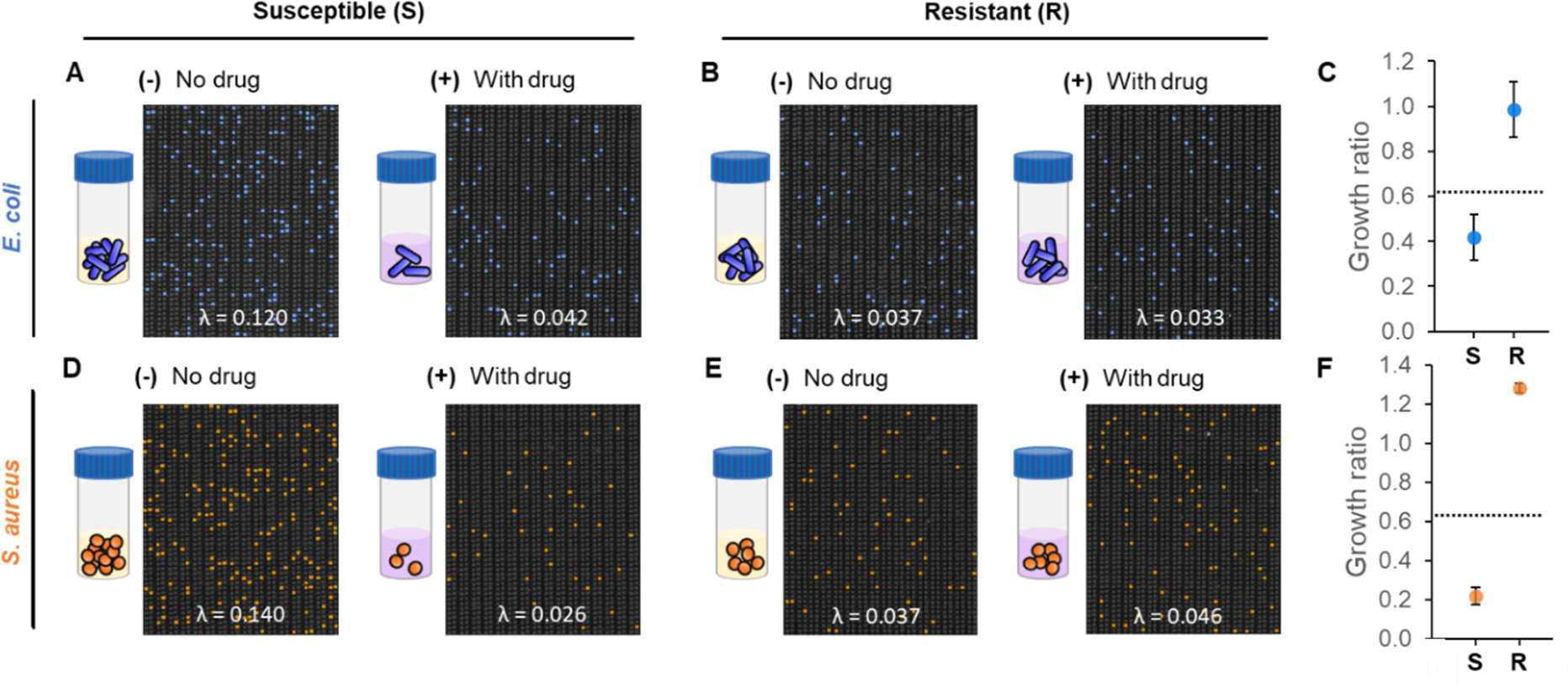
Characterization of pheno-molecular AST with dPCR-HRM-based quantitative analysis in Nanoarray. A panel of 4 bacterial strains with distinct susceptibilities to gentamicin is used to characterize the performance of pheno-molecular AST coupled with dPCR-HRM-based quantitative analysis. In all cases, the gentamicin-treated aliquot and the no-antibiotic control aliquot are incubated for only 30 min and immediately analyzed via dPCR-HRM in the Nanoarray. Here, species-specific, color-coded Nanoarray images acquired from dPCR-HRM provide effective representation of the pheno-molecular AST results. A) For gentamicin-susceptible *E. coli*, significantly more *E. coli* DNA is measured from the no-gentamicin control aliquot (λ = 0.120) than the gentamicin-treated sample aliquot (λ = 0.042). B) For gentamicin-resistant *E. coli*, comparable amounts of *E. coli* DNA are measured from the no-gentamicin control aliquot (λ = 0.037) and the gentamicin-treated sample aliquot (λ = 0.033). C) The “growth ratios” of gentamicin-susceptible *E. coli* and gentamicin-resistant *E. coli* – calculated from the ratio between λ of the gentamicin-treated sample and λ of the no-gentamicin control from duplicate experiments – are 0.418 ± 0.100 and 0.985 ± 0.124, respectively. In the growth ratio plot, the dashed line represents the susceptibility threshold for determining whether the bacteria strain is susceptible (S) or resistant (R) to gentamicin, which is set at 3 standard deviations from the mean of the growth ratio for the gentamicin-resistant *E. coli*. Similar results are observed for *S. aureus*, where D) significantly more *S. aureus* DNA is measured from the no-gentamicin control aliquot (λ = 0.140) than the gentamicin-treated sample aliquot (λ = 0.026) for gentamicin-susceptible *S. aureus*, and E) comparable amounts of *S. aureus* DNA are measured from the no-gentamicin control aliquot (λ = 0.037) and the gentamicin-treated sample aliquot (λ = 0.046) for gentamicin-resistant *S. aureus*. F) The growth ratios from duplicate experiments are 0.217 ± 0.043 for gentamicin-susceptible *S. aureus* and 1.282 ± 0.024 for gentamicin-resistant *S. aureus*, and correctly fall below and above the susceptibility threshold, respectively.

we calculate the “growth ratio” to concisely represent bacterial susceptibility and resistance to drugs. Here, the growth ratios of the gentamicin-susceptible *E. coli* strain and the gentamicin-resistant *E. coli* strain were 0.418 ± 0.100 and 0.985 ± 0.124, respectively (Figure 4C). The clearly distinguishable growth ratios demonstrate that, when coupling pheno-molecular AST with dPCR-HRM in the Nanoarray, even 30 min of exposure was sufficient for completing the AST that effectively ascertained the susceptibility or resistance of *E. coli* strains to gentamicin.

Similarly, our Nanoarray accurately identified the susceptibility profile of *S. aureus* to gentamicin. After 30 min incubation with 0 or 4 μg mL^−1^ of gentamicin, for gentamicin-susceptible *S. aureus*, we measured 0.140 λ and substantially lower 0.026 λ from the no-gentamicin control and the gentamicin-treated sample, respectively, indicating growth inhibition from gentamicin to this susceptible *S. aureus* (Figure 4D). On the contrary, for gentamicin-resistant *S. aureus*, we observed similar 0.037 λ and 0.046 λ from the no-gentamicin control and the gentamicin-treated sample, respectively (Figure 4E). The similar λʼs suggest that gentamicin did not affect the growth of this resistant *S. aureus*. The growth ratio of the gentamicin-susceptible *S. aureus* was 0.217 ± 0.043, whereas the growth ratio of the gentamicin-resistant *S. aureus* was 1.282 ± 0.024 (Figure 4F). The distinct growth ratios between these two strains demonstrate that performing pheno-molecular AST with Nanoarray effectively identified the susceptibility profiles of *S. aureus*. To the best of our knowledge, this is the first demonstration of rapid pheno-molecular AST with *S. aureus* with only 30 min of antibiotic exposure.

Based on the growth ratios, we established a “susceptibility threshold” for distinguishing resistance from susceptibility for subsequent AST experiments. Our experimental results show that gentamicin-resistant strains of *E. coli* and *S. aureus* both had growth ratios of ~1, which reflect bacterial growth even in the presence of gentamicin. Meanwhile, both susceptible strains had growth ratios of < 1, indicating that gentamicin could effectively prevent their growth. Here, we set the susceptibility threshold that is applicable for both species by subtracting 3 standard deviations from the mean of the growth ratio for the gentamicin-resistant *E. coli* and obtained 0.614 as our susceptibility threshold. In subsequent experiments, bacteria samples with growth ratios below the susceptibility threshold will be identified as susceptible. Conversely, bacteria samples with growth ratios above the susceptibility threshold will be called resistant.

### Bacteria ID and Pheno-Molecular AST from Complex Samples

By performing dPCR-HRM in the Nanoarray, our platform offers unique capacity for analyzing complex samples – samples that may contain multiple species of bacteria in a clinically relevant sample matrix – without additional sample processing or species isolation steps. For initial demonstration, we used the Nanoarray to identify and enumerate a polymicrobial sample that contains two bacterial species. This mock polymicrobial sample was generated by mixing *S. aureus* with *P. mirabilis* at equivalent amount. After dPCR-HRM and species identification using our identification algorithm, 0.011 λ were identified as *S. aureus* and relatively similar amount of 0.012 λ were identified as *P. mirabilis* (Figure 5A). This result evidently supports that our Nanoarray is capable of simultaneously identifying multiple bacterial species in a single sample.

**Figure 5.**
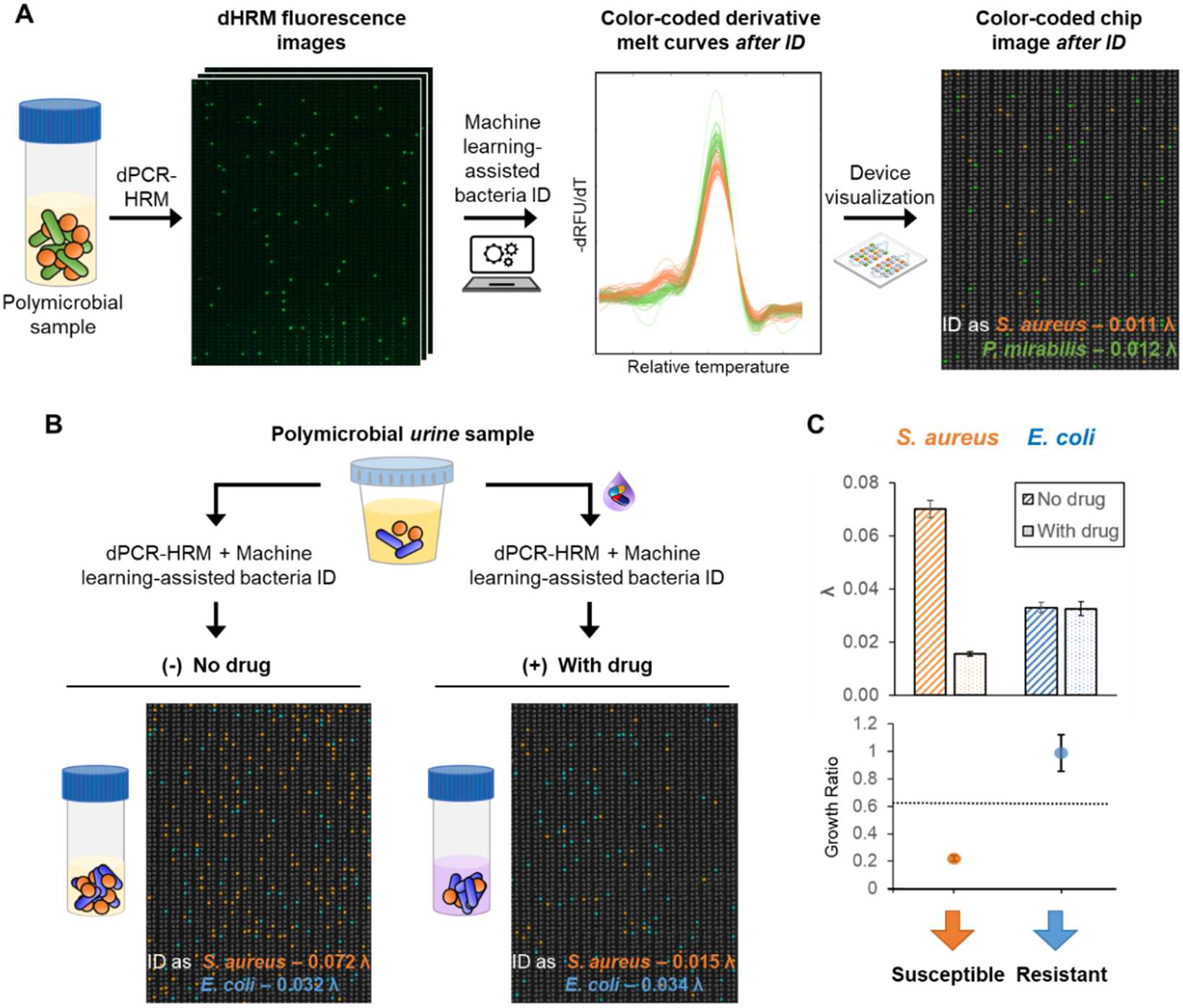
Bacteria ID and pheno-molecular AST from complex samples in Nanoarray. A) Our platform offers unique capability for analyzing polymicrobial samples. For example, equal concentrations of *S. aureus* and *P. mirabilis* spiked in the same sample are both correctly identified and enumerated by performing dPCR-HRM and machine learning-assisted bacteria ID. The simultaneous ID results for both species can be clearly visualized through digital melt curves, as well as the corresponding species-specific, color-coded Nanoarray image. B) Our platform is also capable of achieving comprehensive bacteria ID and pheno-molecular AST for urine sample containing multiple bacteria with distinct antibiotic susceptibilities. Here, samples containing 10% urine with spiked-in gentamicin-susceptible *S. aureus* and gentamicin-resistant *E. coli* are used for testing. Following pheno-molecular AST, dPCR-HRM, and machine learning-assisted bacteria ID, color-coded Nanoarray images clearly show that not only are both species in the sample are identified, but more *S. aureus* DNA is detected in the no-gentamicin control (λ = 0.072 for the no-gentamicin control and λ = 0.015 for the gentamicin-treated sample) and comparable amounts of *E. coli* DNA are detected in the no-gentamicin control (λ = 0.032) and the gentamicin-treated sample (λ = 0.034). C) After multiple tests, the corresponding λʼs and growth ratios confirm the identity and the gentamicin susceptibility for both gentamicin-susceptible *S. aureus* and gentamicin-resistant *E. coli*. These results validate the repeatability of our platform in achieving bacteria ID and AST in polymicrobial urine samples.

As the final demonstration of our method, we identified two species of bacteria and determined their susceptibilities to gentamicin directly from a simulated polymicrobial urine sample. Here, we created the simulated polymicrobial urine sample by spiking both gentamicin-susceptible *S. aureus* and gentamicin-resistant *E. coli* in culture medium laced with 10% culture-negative (*i.e*., bacteria-free) urine sample (Figure 5B). After 30 min gentamicin exposure, in a representative no-gentamicin control, 0.072 λ were identified as *S. aureus* and 0.032 λ were identified as *E. coli*. On the other hand, from the corresponding gentamicin-treated sample, we identified 0.015 λ as *S. aureus* and 0.034 λ as *E. coli* (Figure 5B, representative color-coded Nanoarray devices are shown; *S. aureus* – orange, *E. coli* – blue). Similarly, from duplicate experiments, growth ratios were calculated to be 0.224 ± 0.022 and 0.991 ± 0.133 for *S. aureus* and *E. coli*, respectively (Figure 5C). By comparing these ratios with the susceptibility threshold previously determined for these two bacterial species, *S. aureus* was determined to be gentamicin-susceptible while *E. coli* was determined to be gentamicin-resistant – both in accordance with what we expected. These results demonstrate the use of Nanoarray for simultaneously identifying multiple bacterial species and performing AST, even in the presence of urine.

## Conclusions

We have developed the first microfluidic-based universal dPCR and machine learning-assisted dHRM analysis platform that enables broad bacteria ID and rapid pheno-molecular AST for addressing critical unmet needs in the clinical diagnosis of infectious diseases. We first developed the Nanoarray device and performed dPCR-HRM to broadly detect 5 common UTI species based on their universal 16S rRNA gene and generate hundreds of melt curves for each bacterial species in parallel, which have been stored in our digital melt curve database. We subsequently created our machine learning-based melt curve classification algorithm, and used it in tandem with reference melt curves generated directly in each Nanoarray device to achieve bacteria ID. We have also shown the capability of the Nanoarray in measuring the concentrations of E. coli genomic DNA across three titrations. Precise quantification of bacterial DNA in the Nanoarray allowed us to accurately determine susceptibility profiles of gentamicin-susceptible and gentamicin-resistant strains of E. coli and S. aureus via pheno-molecular AST with as little as 30 min exposure to gentamicin. Moreover, sample digitization in our Nanoarray allows each bacterial DNA molecule to be individually interrogated through dPCR-HRM, which enables polymicrobial detection. Leveraging this unique capability, we analyzed a spiked, polymicrobial urine sample and correctly identified the gentamicin-sensitive E. coli and the gentamicin-resistant S. aureus that were both present in the sample with a turnaround time of ~4 hours. These results illustrate the potential of our platform as a comprehensive and rapid diagnostic method for infectious diseases.

We highlight several technical and conceptual advances of our work. First, by digitizing DNA molecules through limiting dilution in the Nanoarray, our platform ensures that each DNA molecule can be individually quantified and analyzed, and thus enables the detection of individual bacterial species in polymicrobial samples. This advance addresses an intrinsic limitation of PCR-HRM-based bacteria identification performed in bulk (*e.g*., reaction tubes), which has a limited capacity in diagnosing polymicrobial samples because composite melt curves from multiple bacterial species in the sample cannot be easily resolved. Second, we demonstrate that, in the Nanoarray, dPCR-HRM offers comparable level of quantitative precision as dPCR alone while adding the potential benefit of reducing false-positives. Third, within a single Nanoarray device, we can generate hundreds or even thousands of digital melt curves for each species from a single experiment, allowing us to rapidly accumulate large sets of digital melt curves in our database for supervising, training, and improving the robustness of our machine learning-based digital melt curve identification algorithm. We also introduce using a reference module in the Nanoarray to perform dPCR control and generate reference digital melt curves in situ, which provides effective means for mitigating potential variations in digital melt curves due to experimental conditions. Finally, we demonstrate a new approach for achieving digital melt curve-based identification, in which we first classify the melting temperatures and then differentiates the shape of digital melt curves. This approach provides a solid conceptual framework as we expand our bacteria ID capacity.

We also envision making a number of improvements and expanding the scope of testing our platform so that our platform can be used in clinical settings to enable precision-directed therapy, improve patient outcome, and reduce the spread of antibiotic resistance in the future. For example, it is imperative to continue simplifying and accelerating our method. To this end, we can expand the dynamic range of the Nanoarray by increasing the number and/or the volume of the nanowells to facilitate direct analysis of bacterial DNA without dilution. This strategy could be particularly useful for the diagnosis of UTIs, for which the relevant bacteria concentrations range from 10^4^ to 10^7^ colony forming units per mL. In addition, we can implement a rapid PCR assay in the Nanoarray to shorten the turnaround time of our method. Although we have established the conceptual framework for achieving broad bacteria ID, we must continue expanding the number of bacterial species in our digital melt curve database and refining our digital melt curve identification algorithm accordingly to broaden our capacity of bacteria ID. Similarly, for pheno-molecular AST, we must test our platform against bacteria with various growth rates, as well as antibiotics with different mechanisms and at their respective minimum inhibitory concentrations (MICs). The expanded testing conditions would allow us to optimize the incubation time and refine the antibiotic susceptibility threshold for various bacterial species. Finally, our platform offers unique capacity for diagnosing polymicrobial infections, which present an increasingly relevant clinical challenge, especially for UTIs.^[12,17,18]^ For example, up to 1 in 3 elderly UTI patients are polymicrobial^[60]^ and up to 31% of catheter-associated UTIs especially with long-term catheters are polymicrobial.^[10,61]^ We therefore envision testing the performance of our platform in diagnosing polymicrobial infections from urine and potentially other types of clinical samples. Given the demonstrated capability and the potential for expansion, we believe that our platform, when fully developed, will become a useful diagnostic tool for rapid pathogen identification and antibiotic susceptibility testing from complex specimen.

## Acknowledgements

We thank Dr. Kathleen E. Mach and Prof. Joseph C. Liao from Veterans Affairs Palo Alto Health Care System for providing urine specimen, and Aniruddha Kaushik for his help in coordinating. We also thank Dr. Helena C. Zec, Dr. Wei Liu, and Dr. Liben Chen for the initial design of the microfluidic devices and the imaging system. This research is supported by National Institutes of Health (grant R01AI117032, R01AI137272, R01AI138978). K.H. is financially supported through a Hartwell Postdoctoral Fellowship.

## Supporting Information

Materials and methods, supporting figures for thermal-optical setup, Nanoarray device fabrication, benchtop- and Nanoarray-generated melt curves comparison, fluorescent images of all bacterial species for generating digital melt curve database, dHRM-based bacteria identification using Nanoarray digital melt, principal component analysis of digital melt curves from 5 bacterial species, no template control experiment on Nanoarray, schematic of bacterial ID/AST using Nanoarray digital melt, and melt curve acquisition and data processing

