## Supplementary Figure for "Machine Learning-Assisted Digital PCR and Melt Enables Broad Bacteria Identification and Pheno-Molecular Antimicrobial Susceptibility Test"

### Table of Contents

Material and Methods……………………………………………..……………………………...S2
Figure S1. Thermal-optical setup for dHRM analysis…………………………………………………S8
Figure S2. Nanoarray device fabrication………………………………………………………………S9
Figure S3. Benchtop- and Nanoarray-generated melt curves comparison……………………………S10
Figure S4. Fluorescent images of all bacterial species for generating digital melt curve database…..S10
Figure S5. dHRM-based bacteria identification using Nanoarray digital melt……………………….S11
Figure S6. Principal component analysis of digital melt curves from 5 bacterial species……………S12
Figure S7. No template control experiment on Nanoarray……………………………………………S13
Figure S8. Schematic of bacterial ID/AST using Nanoarray digital melt…………………………….S14
Figure S9. Melt curve acquisition and data processing………………………………………………S15
References………………………………………………………………………………………….S16

**Materials and Methods**

*Bacterial Strains and Storage*:
Seven bacterial strains and quantified genomic DNA of *E. coli* used in this study were purchased from the American Type Culture Collection (ATCC; Manassas, VA): *S. agalactiae* ATCC 13813, *P .mirabilis* ATCC 12453, *S. aureus* ATCC 29213, *K. pneumoniae* ATCC BAA-1705, *E. coli* ATCC 25922, *E. coli* ATCC BAA-2471, and *S. aureus* ATCC BAA-44. Of these, *E. coli* BAA-2471 and *S. aureus* BAA-44 are multi-drug resistant. The bacterial stocks received from ATCC were first plated on tryptic soy agar plates (TSA; BD Diagnostics, Sparks, MD) overnight at 37 °C. Subsequently, from each plate, a number of colonies were picked and cultured in tryptic soy broth (TSB; BD Diagnostics, Sparks, MD) at 37 °C. After culturing for ~8 hours, all bacterial cultures were supplemented with sterile glycerol at 20% v/v (Sigma-Aldrich, St. Louis, MO), aliquoted, and kept at -80 °C for long-term storage. Finally, after the aliquots were completely frozen (in ~3 days), at least one aliquot from each bacterial strain was thawed and quantified via plating (in TSA plates) to estimate the stock concentration of each bacterial strain.

*Design of Nanoarray Devices*:
The Nanoarray is composed of 3 separated but identical modules. Each module has a single-inlet and a single-outlet that are designed to interface with blunt-end needles to facilitate sample loading and partitioning. The inlet splits into 28 100-µm-wide branch channels that run in parallel to each other through the module and to the outlet. At the center of the module, each branch channel connects to 180 nanowells. Each module therefore houses a total of 5,040 nanowells. The nanowells are 125 µm in length and 100 µm in width. The branch channels and the nanowells have a uniform height of ~80 µm. Thus, each nanowell is 1 nL in volume, and each Nanoarray module has an analysis volume of ~5 µL. Importantly, the array area of the device is designed to have an "ultra-thin" profile[1] to minimize sample evaporation during thermocycling and to reduce background fluorescence.

*Fabrication of Nanoarray Devices*:
Our Nanoarray devices were made with polydimethylsiloxane (PDMS) and glass via our unique ultra-thin layering technique51 based on soft lithography and PDMS-glass bonding. We began with microfabricating a reusable casting mold by spinning a 80-µm-thick layer of SU-8 3050 photoresist (MicroChem, Westborough, MA) onto a dehydrated and plasmas-treated 4 in. silicon wafer (Polishing Corporation of America, Santa Clara, CA) and patterning the Nanoarray channels and nanowells using standard photolithography. Prior to casting the PDMS fluidic pattern layer of each Nanoarray device, the casting mold was treated with chlorotrimethylsilane (Sigma-Aldrich, St. Louis, MO) via vapor deposition in a vacuum chamber for ~10 min to minimize adhesion of PDMS to the photoresist and the wafer surface. A 15:1 mixture of PDMS (SYLGARD 184 Silicone Elastomer Kits (Dow Corning, Midland, MI)) was then spun on the mold at 500 rpm to fabricate the fluidic pattern layer (~80-µm thick). In parallel, a 6:1 mixture of PDMS was spun on a blank 4 in. silicon wafer at 100 rpm to fabricate a sacrificial PDMS layer (~1-mm thick) whose function is to facilitate peeling the thin fluidic pattern layer off the casting mold without damage. Both PDMS layers were baked at 80 °C for 6 min. The solidified sacrificial PDMS layer was then peeled off the blank wafer, laid over the pattern layer, and baked at 80 °C for 6 min to promote adhesion between the two PDMS layers. In doing so, these two thermally-adhered layers could be jointly peeled off the casting mold, cut into individual

devices, and hole-punched at the inlets and outlets without separation. Next, to fabricate the base of the Nanoarray device, a layer of 15:1 PDMS was spun at 2100 rpm onto a DI water cleaned and blow-dried blank glass slide (75 mm × 50 mm × 1 mm thick; Ted Pella, Redding, CA) and then baked at 80 °C for 6 min. The pattern PDMS layer and the blank PDMS layer on the glass slide were simultaneously treated with oxygen plasma (500 mTorr and 42 W for 45 s), bonded, and baked at 80 °C for 6 min. After baking, the pattern layer became irreversibly bonded to the blank PDMS layer on the glass slide, and the sacrificial PDMS layer could be peeled from the pattern layer. The top of the pattern layer, a glass coverslip (24 mm wide × 60 mm long × 0.13 - 0.16 mm thick; Ted Pella, Redding, CA), and two PDMS tubing adapters (10:1 PDMS, ~4 mm thick, hole-punched) were simultaneously treated with oxygen plasma. The coverslip was then laid over and bonded to the array area of the pattern layer, while the two adapters were laid over and bonded to the inlets and the outlets of the pattern layer. Each completely assembled Nanoarray device was baked at 80 °C overnight. Finally, prior to experimentation, the inlets and outlets of the device were sealed with thin adhesive tapes, and the device was placed in a vacuum chamber for a minimum of 4 hours.

*Overview of Bacteria ID and Pheno-Molecular AST in Nanoarray*:
To perform digital-based bacterial ID/AST in Nanoarrays, we began with sample preparation where a bacterial sample was divided into two portions: one portion was added with the antibiotic of interest, while the other portion was without antibiotics and served as the no drug control. Both portions were then briefly incubated and followed by DNA extraction. Extracted DNA was mixed with PCR mixture and loaded into Nanoarrays to perform dPCR-HRM. Finally, the results were analyzed to achieve bacterial ID and AST (Figure S8). Each of these steps is described in detail following this section.

*Sample Preparation and Antibiotic Exposure*:
For monomicrobial testing in culture broth, frozen stocks of bacteria were thawed and titrated to target concentrations with Muller-Hinton II cation adjusted broth (MHII, BD Diagnostics, Sparks, MD). Gentamicin (Quality Biological Inc., Gaithersburg, MD) was used as the antibiotic of interest and diluted with molecular grade water (Quality Biological Inc., Gaithersburg, MD). The experiments were done in duplicate. The samples were first prepared with MHII containing final concentration of $5\times10^6$ CFU/mL of bacteria before splitting into 2 portions that contained 0 (no drug control) and 4 µg/mL of gentamicin at the final volume of 1 mL. Both portions were then incubated briefly for 30 min at 37 °C.

For polymicrobial testing in urine sample, a boric acid-free, culture-negative urine sample was obtained from the Department of Urology at Stanford University and stored at 4 °C before testing. The urine sample was diluted 10× in Mueller-Hinton II broth (MHII; BD Diagnostics, Sparks, MD) and added with gentamicin-susceptible *S. aureus* and gentamicin-resistant *E. coli* at $5\times10^7$ CFU/mL and $1.25\times10^6$ CFU/mL respectively to mimic a polymicrobial infection. The experiment was conducted in duplicate. We started by splitting the mixed sample into two reactions of 1 mL each to incubate with and without 4 µg/mL of gentamicin (Quality Biological Inc., Gaithersburg, MD) for 30 min, followed by DNA extraction prior to dPCR-HRM on Nanoarrays.

*Bacterial DNA Extraction*:

After incubation, each bacterial culture aliquot was spun at 10,000× g for 10 min to discard broth supernatant, washed with 500 µL of molecular grade water, and spun again at 10,000× g for 10 min to discard water supernatant. To perform DNA extraction, 100 µL of QuickExtract™ DNA Extraction Solution (Lucigen, Madison, WI) and 1000 U of Ready-Lyse™ Lysozyme Solution (Lucigen, Madison, WI) were added into each pellet followed by incubating at room temperature for 1 hour. The concentration of the obtained DNA was optionally estimated by performing real-time PCR with titrations using a CFX96 Touch Real-time PCR Detection System (Bio-Rad, Hercules, CA) with CFX Manager software. Finally, the extracted DNA was then diluted 25-fold or higher to obtain desired concentrations before adding into PCR mixes to perform dPCR-HRM on Nanoarrays.

*Preparation of Universal Digital PCR Mixture*:
Typically, 3 universal digital PCR reactions were prepared for the 3 modules of the Nanoarray. One of the reactions is for performing dPCR-HRM with *E. coli* quantified genomic DNA to attain *in situ* dPCR-HRM control and reference digital melt curves. The other two reactions are for testing the experimental conditions of interest (*e.g.*, drug testing sample and no drug control, or two bacterial species).

Our universal digital PCR mixture is composed of 2 µL diluted bacterial DNA, 1× Gold Buffer (Thermo Fisher Scientific, Waltham, MA), 3.5 mM MgCl2 (Thermo Fisher Scientific, Waltham, MA), 1× Evagreen (Biotium, Freemont, CA), 1× ROX (Thermo Fisher Scientific, Waltham, MA), 1 mg/mL BSA (New England Biolabs, Ipswich, MA), 0.01% Tween 20 (Sigma-Aldrich, St. Louis, MO), 200 µM of each deoxynucleotide triphosphate (Thermo Fisher Scientific, Waltham, MA), 0.3 µM forward primer (5'-GYGGCGNACGGGTGAGTAA-3'; Integrated DNA Technologies, Coralville, IA), 0.3 µM reverse primer (5'-AGCTGACGACANCCATGCA-3'; Integrated DNA Technologies, Coralville, IA), 0.025 U/µL of Amplitaq Gold LD (Thermo Fisher Scientific, Waltham, MA), and Ultra Pure PCR water (Quality Biological Inc., Gaithersburg, MD) to bring the final reaction volume to 40 µL. The amplified products from our PCR reaction were ~970 bp in length.

*PCR Mixture Loading and Digitization in Nanoarray*:
In this work, loading and digitization of PCR mixtures in Nanoarrays were achieved via vacuum-assisted loading and oil-driven digitization.[1,2] Specifically, the PCR mixes were first drawn into blunt-end needles and 1-ft-long sections of Tygon tubing (0.02” inner diameter, Cole-Parmer, Vernon Hills, IL) with empty 1-mL syringes. Then, immediately upon taking the Nanoarrays out of the vacuum chamber, the needles of the tubing were used to puncture through the adhesive tape covering the inlets of Nanoarrays and initiate sample loading. Due to the desiccation, negative pressure within the devices allowed the PCR mixture to fill the entire device and the nanowells. A partitioning fluid was prepared by mixing 5 g 100 cSt silicone oil (Sigma-Aldrich, St. Louis, MO) with 1 g PDMS (10:1), and drawn into blunt-end needles and 4-ft-long sections of Tygon tubings. Once the PCR mixture had completely filled all nanowells, the PCR mixture tubings were removed from the inlets, and the partitioning fluid tubings were immediately inserted into device inlets and pressurized at ~14 psi using a pressure regulator. After removing adhesive tape covering the device outlets, the partitioning fluid flew through branch channels of the devices without entering nanowells, thereby isolating and digitizing all nanowells within the Nanoarrays and readying the devices for dPCR-HRM.

*dPCR-HRM in Nanoarray*:
After loading and digitization of PCR mixtures, the Nanoarrays were first placed on a flatbed thermal cycler (Proflex, Thermo Fisher Scientific, Waltham, MA) to perform dPCR with the following cycling conditions: a hot start at 95 °C for 10 min, followed by 60 cycles of 95 °C for 15 s, 67 °C for 15 s, and 72 °C for 60 s, and a final extension at 72 °C for 7 min. The partitioning fluid was pressurized at ~14 psi throughout dPCR. Notably, due to the addition of the PDMS component, the partitioning fluid would solidify over the course of the PCR reaction, producing permanent barriers that prevented PCR products from leaking out of nanowells even in the absence of further pressurization. Thus, once dPCR was completed, the Nanoarrays were disconnected from the partitioning fluid tubing (and hence the pressure regulator), removed from the flatbed thermal cycler, and taken to the digital melt instrument to perform dHRM. The digital melt instrument was composed of a flatbed heater, a blue LED array (ThorLabs, Newton, NJ) for fluorescence excitation, and a Sony ILCE camera with a 50 mm lens (Canon, Melville, NY) coupled with a green emission filter (Omega Optical, Brattleboro, VT) for fluorescence image acquisition. Nanoarray devices were secured with electrical tape on a blank silicon wafer (Polishing Corporation of America, Santa Clara, CA), which was placed on the flatbed heater of the digital melt instrument to reduce background fluorescence that can arise from light scattering. To perform dHRM, the temperature was ramped from 75 to 99 °C at 0.1 °C per 2 s increments and the Nanoarrays were illuminated by the LED array, while concurrently, fluorescence images were captured at 1 Hz with the camera, resulting in 480 temperature-lapse fluorescence images for analysis.

*Image Analysis and Melt Curve Extraction*:
Temperature-lapse fluorescence images from each module of the Nanoarray were converted into digital melt curves using a custom-developed melt curve extraction program written in MATLAB. All 480 fluorescence images were first converted from RGB to grayscale and then aligned to the first image, which was acquired at 75 °C, to ensure that images that might have drifted due to unwanted movement of the Nanoarray device during HRM would be corrected. A grid mask that specified the locations of all nanowells from the Nanoarray module was then created from the grid mask template originally created based on the device design. Specifically, first, the locations of the four corners of the first fluorescence image were input by users to specify the coordinates of the four corners of the grid mask (Figure S9). The grid mask was then homography transformed (*i.e.,* stretched and tilted) to precisely map the coordinates of the grid mask with the locations of the nanowells. Finally, the center of the grid mask over each nanowell was moved around a 2-by-2 pixel neighborhood to ensure that the grid mask would enclose maximum fluorescence intensity from the nanowell. Using the final grid mask with optimized locations, average fluorescence intensities from all nanowells in the Nanoarray module were measured from all 480 temperature-lapse fluorescence images. Digital melt curves were subsequently generated by binning fluorescence intensities into 0.2 °C increments and plotting the fluorescence intensities as a function of temperature. The Savitzky-Golay filter was applied to preserve the shape of the melt curves while removing noise. Then, negative derivative melt curves (*i.e.*, -dRFU/dT; hereafter referred to as "digital melt curves") were calculated and further smoothed with Savitzky-Golay filter. This image analysis and melt curve extraction processes were repeated for the two other modules of the Nanoarray, resulting in 3 independent groups of digital melt curves.

*dHRM-Based Bacteria ID through Automated Machine Learning Algorithm*:
Using our platform, 320 digital melt curves of 5 bacterial species were first experimentally generated and stored in the digital melt curve database. Subsequently, newly generated digital melt curves of unknown bacterial species from subsequent experiments were analyzed with our custom-developed identification algorithm written in MATLAB to achieve bacteria ID. The automated identification algorithm encompasses 3 main steps: 1) classification of digital melt curves based on *Tm*, 2) normalization and alignment of digital melt curves, and 3) identification of digital melt curves based on comparison with the digital melt curves in the database using our ovoSVM algorithm.[3]

In this work, we classified the digital melt curves of the 5 bacterial species into either the low *Tm* group or the high *Tm* group based on their *Tm* relative to the *Tm* of the reference digital melt curves that were acquired from *E. coli* genomic DNA *in situ*. The classification process began by combining both the digital melt curves from the analyzing module with the *E. coli* reference digital melt curves into a single group and plotting a histogram of the *Tm*. If the *Tm* histogram displayed a single peak, indicating that the digital melt curves from the analyzing module had similar *Tm* as the *E. coli* reference digital melt curves, the digital melt curves were classified into high *Tm* group. On the other hand, if the *Tm* histogram also displayed a second, lower peak, indicating that many digital melt curves from the analyzing module had lower *Tm* than the *E. coli* reference digital melt curves, a *Tm* threshold was then set to the *Tm* at which the local minimum in frequency between the two peaks was found. Then, individual digital melt curve from each nanowell was classified into either the low or the high *Tm* group based on its *Tm* with respect to the *Tm* threshold. Of note, digital melt curves with the same *Tm* as the *Tm* threshold, which could occasionally arise from DNA of mixed species in the nanowells, were classified as unidentified and discarded from further analysis.

Once the digital melt curves from each analyzing module were classified in the appropriate *Tm* group, the shapes of the digital melt curves were used toward identification. To this end, the area under all digital melt curves was first normalized so that potential variations in fluorescence intensities in different nanowells were made equal. To accentuate the differences in the shapes of the digital melt curves, all digital melt curves were aligned at a single point, which was empirically set at the point when -dRFU/dT of the falling edge of the digital melt curve (*i.e.*, past the peak) equaled to 40.

Finally, the digital melt curves from each analyzing module were compared with the digital melt curves in the database using our ovoSVM algorithm to achieve bacteria ID. In our algorithm, each digital melt curve from the analyzing module was pitted against the digital melt curves from two species in a binary comparison to determine which species the digital melt curve of interest was more similar. This was accomplished by comparing the data points from the digital melt curves through a combination of linear kernel function and least squares method in our ovoSVM algorithm. The times that the digital melt curve was classified to each species were scored. Finally, bacterial species was identified as the species with the highest matching score from ovoSVM results. However, bacterial species with less than 0.2% positive wells (~10 out of 5,000 wells) were discarded from the analysis. Additionally, a "difference score" that quantifies the similarity between each newly identified digital melt curve and the digital melt curves of the particular species in the database was calculated. This difference score was obtained by summing the absolute differences between the newly identified digital melt curve and the averaged digital melt curves of the particular species in the database. As such, a small difference score represents a good match between the newly identified digital melt curve and the digital melt curves of the particular species in the database, and thus good identification outcome. In this work, we classified digital melt curves with difference scores above 40 (indicative of a poor match between the digital melt curves) as “unidentified” or “false-positive” even if they were initially identified as a particular species. This strategy thus also offered a ready means for classifying digital melt curves that would arise from bacterial species outside of our database.

*Safety Statement:*
No unexpected or unusually high safety hazards were encountered in this work.

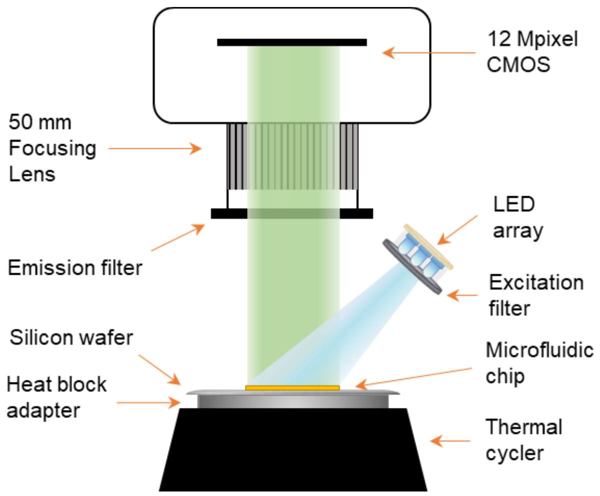


**Figure S1.** Thermal-optical setup for dHRM analysis. A blue LED array with filter was used as a light source to illuminate the microfluidic devices at ~45-degree angle. The device was placed on top of a blank silicon wafer and secured with tape on a commercial flatbed thermocycler. A commercial 12Mpixel CMOS MIL camera with focusing lens and a filter were used to acquire fluorescence images during temperature ramping process in HRM.

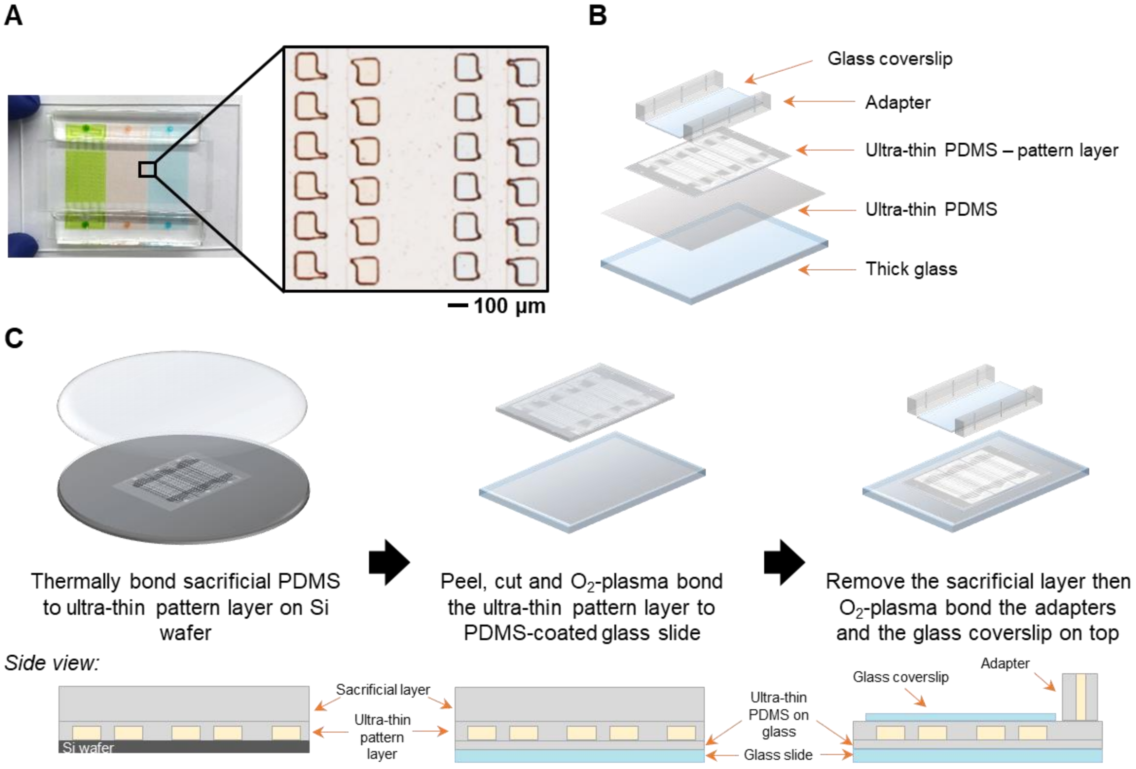


**Figure S2.** Nanoarray device fabrication. (A) A Nanoarray device loaded with 3 different colors of food dye in 3 independent modules. The zoom-in image showed food dye-loaded nanowells after isolating with partitioning fluid. (B) The drawing shows components of a Nanoarray, which comprise of an ultra-thin PDMS-coated thick glass slide for the bottom layer, an ultra-thin PDMS pattern layer, and a glass coverslip between adapters for inlets and outlets. (C) Our fabrication process starts by placing a PDMS sacrificial layer on top of the cured ultra-thin pattern layer on Si wafer, followed by a brief baking step. Then, the temporarily-joint PDMS layers are peeled off from the mold wafer without tearing, cut and $O_2$-plasma bonded to an ultra-thin-PDMS-coated glass slide. After brief baking step, the sacrificial layer is removed before placing adapters and a glass coverslip on the top of the pattern layer via $O_2$-plasma treatment.

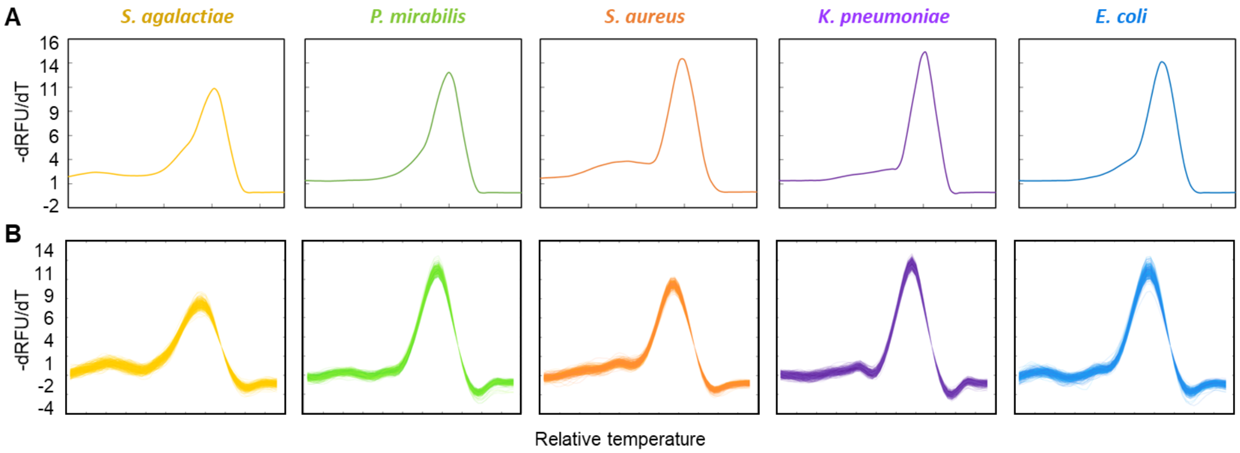


**Figure S3.** Benchtop- and Nanoarray-generated melt curves comparison. (A) The plots show aligned bulk melt curves of 5 bacterial species generated from benchtop instrument. (B) Aligned digital melt curves of 5 bacterial species generating using Nanoarray devices are shown.

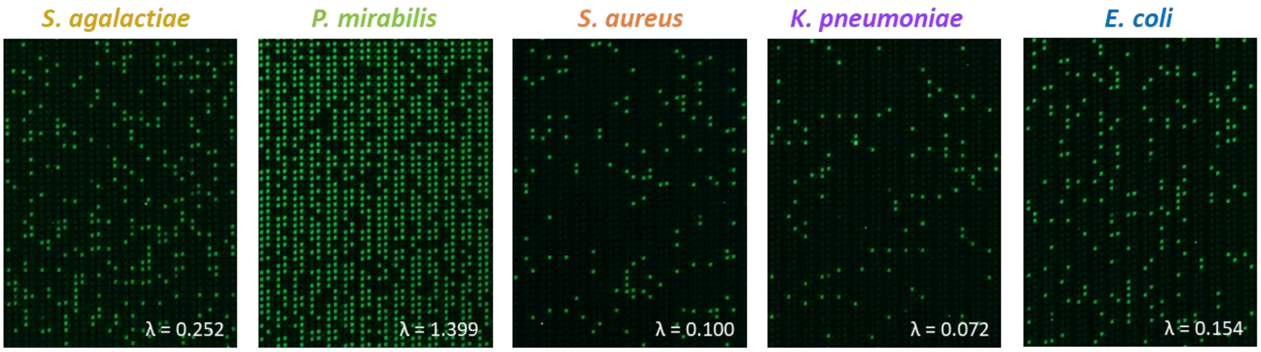


**Figure S4.** Fluorescent images of all bacterial species for generating digital melt curve database. After dPCR, fluorescence images of 5 bacterial species are taken, illustrating successful dPCR amplification in bright green nanowells.

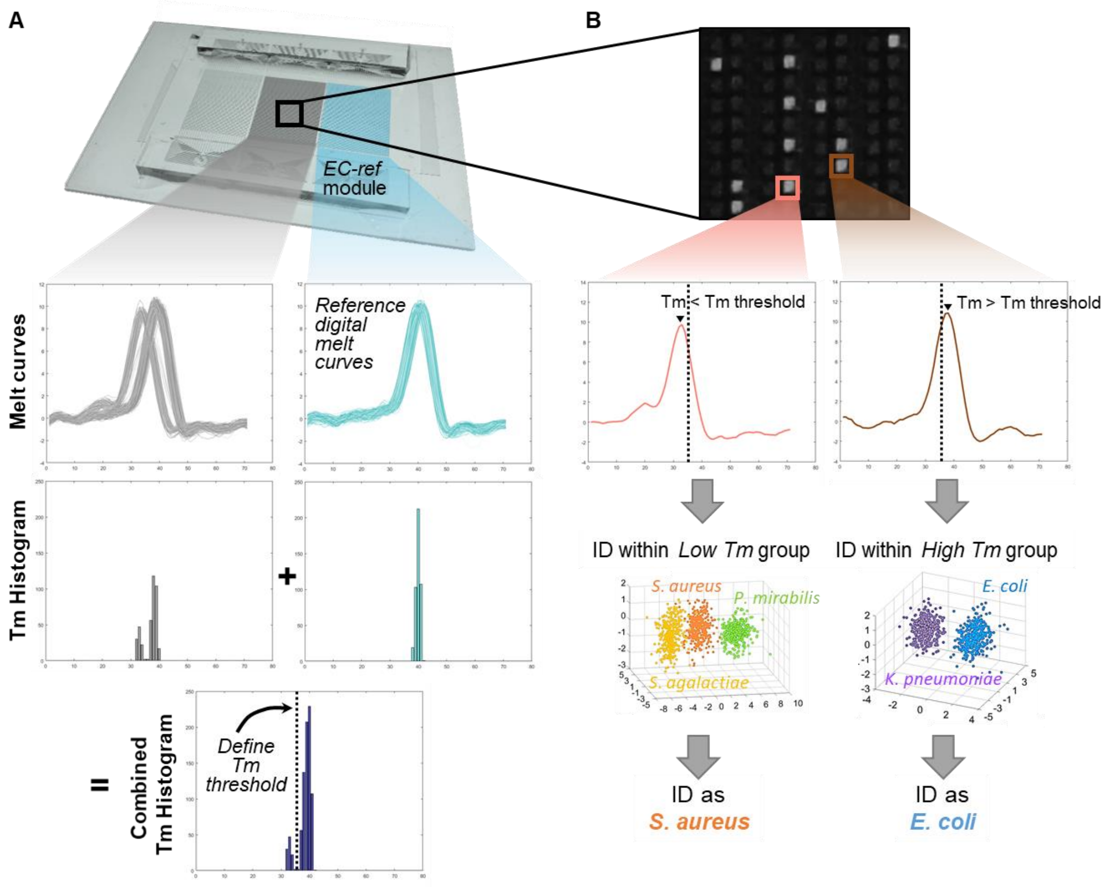


**Figure S5.** dHRM-based bacteria identification using Nanoarray digital melt. (A) To identify bacterial species of unknown melt curves, Tm histograms of the unknown module (gray) and E. coli reference (EC-ref) module (light blue) were summed to produce a combined Tm histogram. In the case that the combined Tm histogram showed two separated peaks, a Tm threshold was defined as the lowest point between those two peaks. (B) Individual melt curve in the unknown module was then categorized into low or high Tm subgroups based on its Tm compared with the defined Tm threshold. Finally, ovoSVM was performed to identify bacterial species of the unknown melt curve by comparing the melt curves with the stored digital melt curves database within the subgroup.

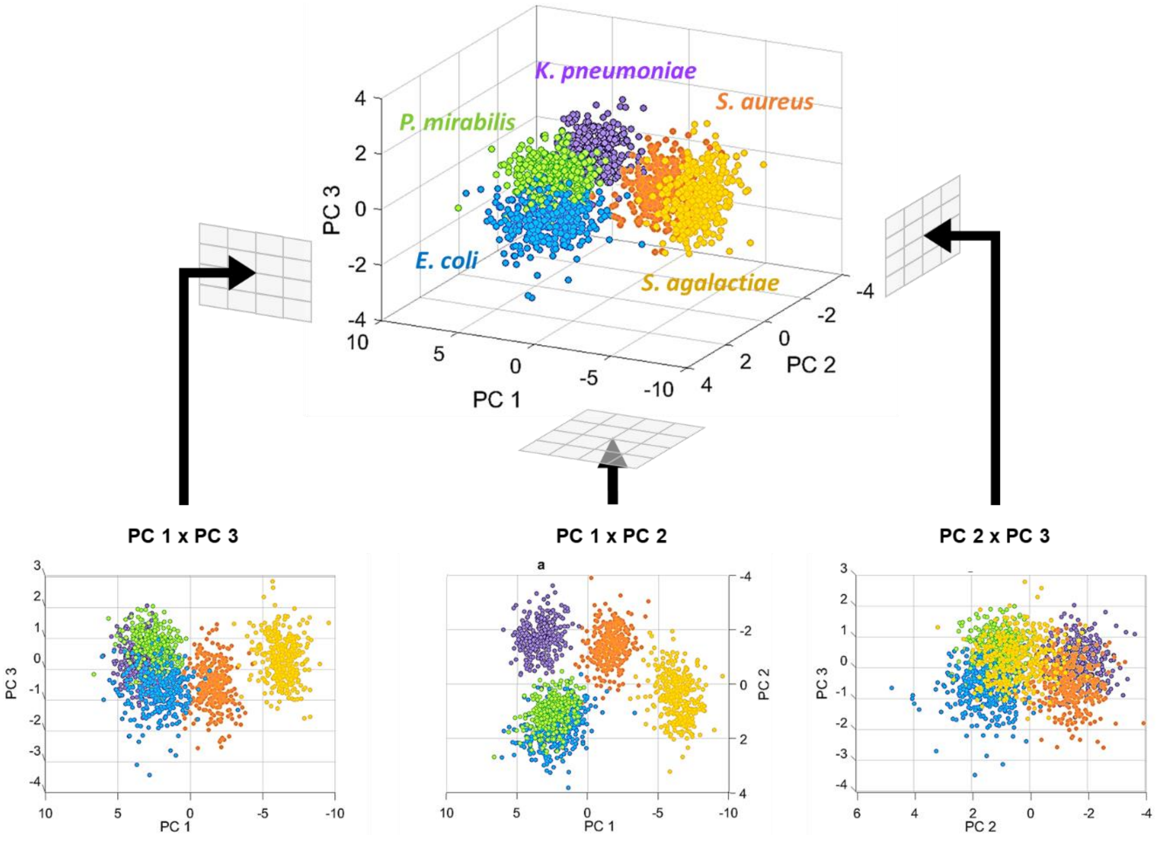


**Figure S6.** Principal component analysis of digital melt curves from 5 bacterial species. The 3D scatterplot shows the first three principal components (PC) of all melt curves (320 melt curves each species) in the digital melt curve library, which explain 90.95% of all variability. Two populations; *P. mirabilis* (green) and *E. coli* (blue) were undistinguishable in any views. Therefore, solely shape comparison after melt curve alignment cannot be used to classify these 5 bacterial species.

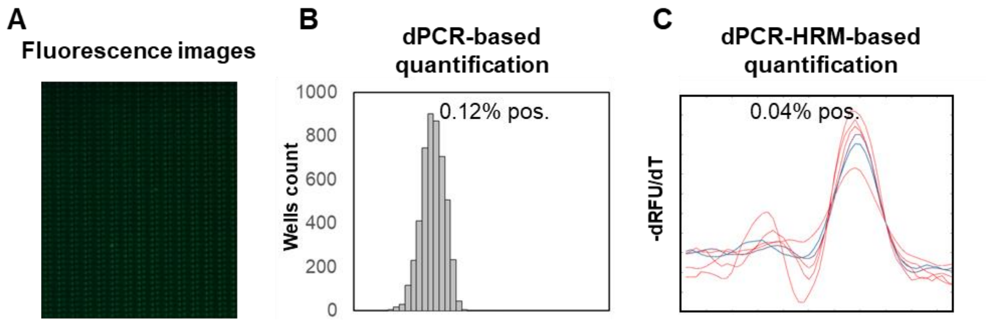


**Figure S7.** No template control experiment on Nanoarray. (A) Fluorescence images of no template module in a Nanoarray device after PCR is shown. (B) The corresponding histogram of fluorescence intensities after PCR revealed that 6 nanowells were positive amplifications (equivalent to 0.12%). (C) Color-coded melt curves are shown after dPCR-HRM-based species identification (4 Red – false-positive, 1 Blue – *E. coli*, and 1 Purple – *K. pneumoniae*). The number of positive reactions became 0.04% after bacteria-ID.

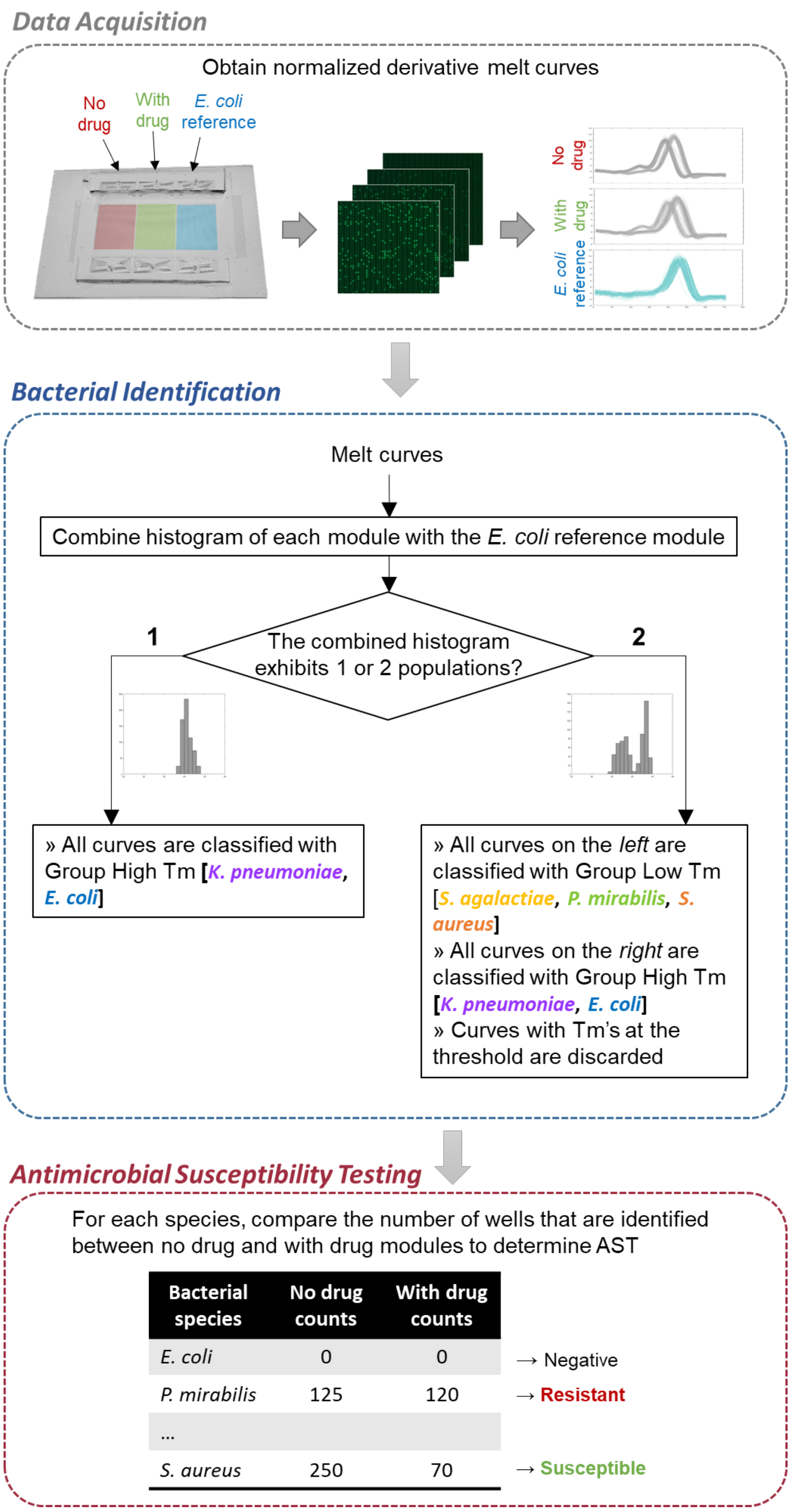


**Figure S8.** Schematic of bacterial ID/AST using Nanoarray digital melt. DNA samples are first loaded into a Nanoarray device to perform dPCR-HRM. Digital melt curves are obtained from temperature-lapse fluorescence images. We then identify bacterial species based on digital melt curves using machine-learning assisted-algorithm. AST profile of each species is determined based on molecular counts after ID.

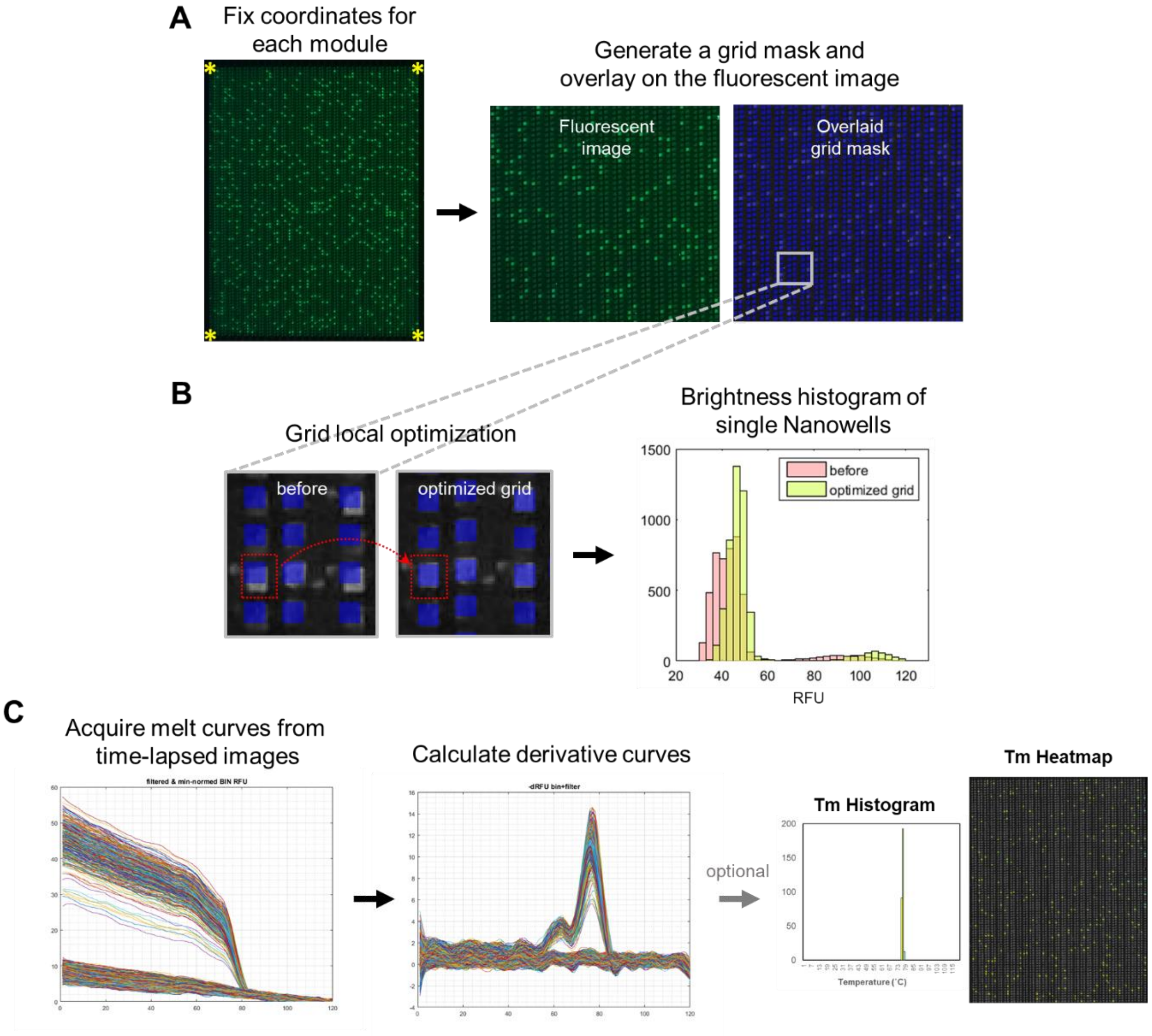


**Figure S9.** Melt curve acquisition and data processing. (A) After obtaining fluorescence images of an entire Nanoarray at each temperature increment, four corner points of the analyzing module were manually selected by users in order to generate a grid mask based on the chip design. For extracting fluorescence values of each nanowell, the grid was homography transformed and (B) performed brightness local optimization over 2-by-2 neighborhood pixels to improve grid-image alignment. The better alignment was observed as the histogram showed increased brightness of each nanowell. In digital dilution, 2 populations of brightness levels were observed; high fluorescence level from positive wells and low fluorescence level from negative or no-target containing wells. (C) The averaged fluorescence level of each well is filtered and plotted against temperature to produce a melt curve. Negative derivative melt curves were calculated. Tm of each curve was defined as the temperature at the peak of each derivative melt curve. A Tm histogram and a Tm heatmap could optionally be generated from melt curves of the entire module.
